# Extending the integrate-and-fire model to account for metabolic dependencies

**DOI:** 10.1101/2020.11.04.367102

**Authors:** Ismael Jaras, Taiki Harada, Marcos E. Orchard, Pedro E. Maldonado, Rodrigo C. Vergara

## Abstract

It is widely accepted that the brain, like any other physical system, is subjected to physical constraints restricting its operation. The brain’s metabolic demands are particularly critical for proper neuronal function, but the impact of these constraints is still poorly understood. Detailed single-neuron models are recently integrating metabolic constraints, but the computational resources these models need, make it difficult to explore the dynamics of extended neural networks imposed by such constraints. Thus, there is a need for a simple-enough neuron model that incorporates metabolic activity and allows us to explore neural network dynamics. This work introduces an energy-dependent leaky integrate-and-fire (LIF) neuronal model extension to account for the effects of metabolic constraints on the single-neuron behavior (EDLIF). This simple energy-dependent model shows better performance predicting real spikes trains -in *spike coincidence* measure sense-than the classical leaky integrate-and-fire model. It can describe the relationship between the average firing rate and the ATP cost, and replicate a neuron’s behavior under a clinical setting such as amyotrophic lateral sclerosis. The simplicity of the energy-dependent model presented here, makes it computationally efficient and thus, suitable to study the dynamics of large neural networks.

**Author summary:** Any physical system or biological tissue is restricted by physical constraints bounding their behavior, and the brain is not free from these constraints. Energetic disorders in the brain have been linked to several neurodegenerative diseases, highlighting the relevance of maintaining a critical balance between energy production and consumption in neurons. These observations motivate the development of mathematical tools that can help to understand the dependence of the brain’s behavior in metabolism. One of the essential building blocks to achieve this task is the mathematical representation of neurons through models, allowing computational simulations of single-neurons and neural networks. Here we construct a simple and computational cheap energy-dependent neuron model that allows the study of neuron’s behavior under an energetic perspective. The introduced neuron model is contrasted with one of the widest-used neuron models and shows better prediction capabilities when real neuron recordings are used. Our model is suitable for replicating neuron’s behavior under a specific neurodegenerative disease, which cannot be achieved by the abovementioned popular model. Our simple model is promising because it allows the simulation and study of neuronal networks under a metabolic-dependent perspective.

## Introduction

The brain consumes a disproportionate amount of energy relative to its mass; in humans, the energy used by this organ is 20% of all the oxygen consumed by a body at rest, while its mass only represents 2% of the total body mass [9,10] [25]. This consumption is not only high but fast, as it consumes energy at a rate ten times faster than the rest of the body [36]. Given these conditions, it is not surprising that energy supply must compensate for the energy expenditure incurred by neurons efficiently to maintain proper brain function [11]. Limitations in energy production have been associated with various neurodegenerative diseases, such as amyotrophic lateral sclerosis (ALS), Leigh, Alzheimer’s, and Parkinson’s disease [12–17]. Together, these arguments present robust evidence that supports the importance of energy administration for the brain’s proper function, motivating the development of modeling strategies accounting for metabolic-dependent dynamics.

During the last decade, some of the classical single-neuron models have been extended to include the relationship between available energy and electrophysiological activity [7,21,26]. For example, the study presented in [21] illustrates how different initial ATP levels affect the behavior of the neuron, leading to different types of activities. Whereas in [7] the Hodgkin-Huxley model was extended to include behavioral dependencies on the neuronal metabolic dynamics to study the effects of energetic neuronal dysfunction. These results support the idea that the neuron’s available energy generates a significant impact on its activity. However, these multi-compartment and highly detailed Hodgkin-Huxley models demand increasing computational resources. This high computational cost makes them impractical for studying the effect of energetic dynamics on large neuronal population networks (for a further review of biological plausibility and computational resources trade-off of different neuronal models, the reader is referred to [22]).

As an alternative, the leaky integrate-and-fire (LIF) model is a popular and widely used single-neuron model characterized for his simplicity while at the same time being sufficiently complex to capture many of the essential features of neural dynamics [6]. These attributes made LIF a suitable tool for the study of large neuronal population dynamics simulations. Despite the popularity and simplicity of the LIF model, it is not clear how we can use it to study large neuronal populations under a metabolic-dependent perspective. The LIF model’s main advantage is its balance between simplicity and capacity to capture essential features of neuronal behavior. Despite these attributes, the model, as presented before -and their consecutive extensions-neglect metabolic-dependence in their dynamic. The main reasons for this omission are (1) that the model assumes that the neuron has unlimited access to energy resources, (2) the available energy is restored instantaneously, and (3) the assumption that, under normal condition, the metabolic effect in the neuronal behavior is negligible.

Notwithstanding, a metabolic imbalance can have a meaningful impact on neuronal behavior and in the development of neurodegenerative diseases. Thus, given the relevance of metabolic dynamic for the brain’s proper function, it is beneficial to have a simple single-neuron model that can account for the metabolic dynamic and neuronal behavior relationship. The model mentioned above allows us to reinterpret the single-neuron under metabolic constraint. In this work, we sought to develop a simple single-neuron model that incorporates metabolic rules. Considering that single-neuron models are one of the essential building blocks for in *silico* extensive neuronal population simulations, this would be the first step required to explore neural network dynamics constrained by metabolic demands. Concretely, we extend the LIF model to account for metabolic-dependencies. Thus, granting the simplicity and computational inexpensiveness but allowing it to describe metabolic dynamics and dependencies. We focused on evaluating two relevant aspects of this single-neuron extended model; (1) compared its performance against LIF, and (2) assess whether metabolic predictions of more complex models can be replicated.

## Materials and methods

We employed the LIF model’s structure, extending it to account for the relationship between energetic dynamics and neuronal activity. We termed this new model the energy-dependent leaky integrate-and-fire model (EDLIF). Specifically, we used an incomplete repolarization mechanism [5] and made it ATP-sensitive, thus making the -post reset- membrane potential ATP-dependent. Given that our derivation of the EDLIF model is strongly inspired by the LIF model’s structure, we started recalling the main aspects of the LIF model. Then, we show how it can be extended through ATP-sensitive incomplete repolarization mechanism. In the development of these concepts, we also analyzed the cost and dynamic of energy consumption in the neuron.

### Leaky Integrate-and-Fire model

The LIF neuron model is a single compartment model that describes the membrane potential in terms of the synaptic inputs and the injected current that it receives. An action potential is generated when the membrane potential reaches a -fixed- threshold (*V_th_*) [6]. The model is described by the dynamics of the neuron’s membrane potential *v_m_*(*t*) in Eq 1:

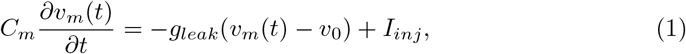

where *C_m_* is the membrane capacitance, *g_leak_* is the conductance associated to the current leakage through the cellular membrane and the passive membrane time constant is 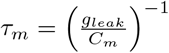. Whereas *I_inj_* describes the injected current to the neuron, it accounts for the effect of synaptic input and the current stimuli by an intracellular electrode.

After the occurrence of an action potential (*i.e.* when *v_m_*(*t*) ≥ *V_th_*) the membrane voltage *v_m_* is resorted to a reset value (*V_reset_*) and a minimum time -called the refractory time-as to pass before a new action potential can be generated. This straightforward conceptualization of the neuron is the root of a wide variety of models extensions aiming to improve the characterization of real neuronal behavior made by the original LIF model [22]. In the next subsection, we will extend the LIF model to introduce the Energy Dependent Leaky Integrate-and-Fire model.

### Energy Dependent Leaky Integrate-and-Fire model

Concerning the limitations of metabolic-dependencies in the LIF model, in this section, we introduce our Energy Dependent Leaky Integrate-and-Fire (EDLIF) model. Our proposed model is, as his name declares, an energy-dependent neuronal model inspired by the LIF. The model’s main objective is to include energy dependence in the neuronal dynamics but maintaining the simplicity of the LIF model, so it requires low computation resources and is suitable for the simulation and study of networks of thousands of neurons through an energy-dependent perspective. In this regard, the dynamic of the membrane potential of neurons will be described in the same way as Eq 1, but the repolarization mechanism will include an energy-dependent relationship, affecting the temporal evolution of the neuron if an energy imbalance is present. In EDLIF the neuron’s energy dynamic will be described by the dynamic of the intracellular ATP concentration.

### ATP dynamics

In our model, ATP dynamics will be characterized considering two collections of processes: those that supply ATP to the neuron (*A_s_*(*t*)), and those that consume ATP (*A_c_*(*t*)). The ATP dynamic concentration (*A*(*t*)) can be formalized by Eq 2 [8]:

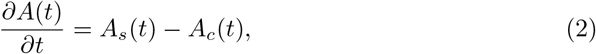

where both terms *A_s_*(*t*) and *A_c_*(*t*) are divided per volume unit.

The observation that the neuron has relatively constant ATP can be interpreted as reflecting homeostatic feedback. Thus, the neuron’s homeostatic mechanisms keep ATP levels close to the homeostatic ATP level (*A_H_*). In this work, the homeostatic feedback will be introduced explicitly in the ATP production dynamics, mathematically formalized by Eq 3. Thus, ATP production depends on two terms: the actual ATP level respect to a homeostatic one (*K*(*A_H_* – *A*(*t*))) and a basal production term (AB) accounting for resting potential and housekeeping activities:

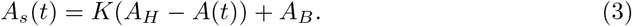

### Energy consumption on neuron

The neuron, like any other cell, requires energy to carry out its activities, which can be classified as signaling and non-signaling [1]. Within the non-signaling activities, the neuron uses energy to carry out maintenance processes (such as organelle traffic and the synthesis of proteins and molecules) and conserve the membrane potential during rest. Signaling activities are present only during the communication periods, such as action and postsynaptic potentials and presynaptic activity associated with synaptic transmission.

Since we are working in the single-neuron model, we were mainly concerned about quantifying the energy expenditure related to general homeostasis maintenance (house-keeping), conditions required for signaling (maintaining resting potential), and signaling (action potentials). Postsynaptic potential and presynaptic activity associated are out of the scope of this study. These three energetic budget items are already estimated, as shown in Table 1. These specific values emerge from combining anatomic and physiologic measurements and theoretical calculation of bottom-up energy budget using biophysical properties in conjunction with electrophysiological and morphological data [25,27].

**Table 1.** Energy consumption. Energy consumption related to different processes in a rat excitatory neuron, using experimental data from [25].

| Activity | Unit energy consumption |
| --- | --- |
| House-Keeping ($E_{HK}$) | $8.41 \times 10^8 \text{ ATP}/(\text{neuron} \cdot s)$ |
| Resting potential ($E_{RP}$) | $3.42 \times 10^8 \text{ ATP}/(\text{neuron} \cdot s)$ |
| Action Potential ($E_{AP}$) | $1.25 \times 10^8 \text{ ATP}/(\text{neuron} \cdot \text{spike})$ |

Because the total amount of energy consumes per action potential is not instantly expended, the consumption is described in a biologically plausible way through the inclusion of a time-varying function (Eq 4) [23]. This smooth decay in time reflects the characteristic dynamic of the metabolic energy expenditure in action potential in time.

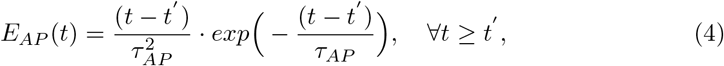

where *t*’ is the time in *ms* at which the action potential occurs and *τ_AP_* = 3945.66 *ms.* See Supporting information for a detailed description of how *τ_AP_* is determined.

### Energy dependencies and incomplete repolarization

So far, we have introduced energy expenditure to EDLIF; however, we still have to include how energy availability may impact neuron physiology. ATP imbalance in the neuron is known to alter sodium-potassium pump. Specifically, a decrease in ATP concentration implies a partial restoration of the resting potential, as it could be inferred from Eq 5 [7]:

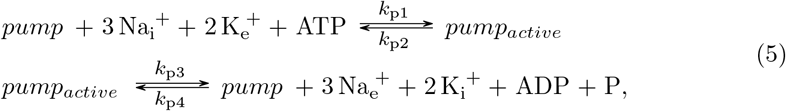

where *‘pump’* and *‘pump_active_’* represents two possible stated for the sodium-potassium pump, 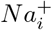 and 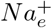 are the intracellular and extracellular sodium concentration, respectively 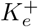 and 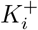 the extracellular and intracellular potassium concentration, respectively and ATP is ATP concentration. This scheme shows that lower ATP concentration disables sodium-potassium pump functioning, reducing the rate at which sodium and potasium concentrations are restored. As such, maintaining the neuron above the resting potential. From Eq 5 follows that, when occurring an action potential and the neuron has ATP deficiency, the neuron will remain slightly depolarized with respect to its resting potential, *i.e.* experiencing a partial repolarization.

One mechanism which allows modifying the membrane potential after the occurrence of an action potential is partial reset. Incomplete repolarization -or partial reset- is a simple and powerful tool for controlling the irregularity of spike trains fired by a leaky integrator neuron model. Previous works have shown that incorporating this mechanism into the LIF model is proven to produce highly irregular firing similar to the one observed in biological neurons [5]. The mechanism works by resetting the potential of the capacitor to *V*(*t*) = *βV_th_* after an action potential occurs, where *V_th_* is the firing threshold and *β* is called the Temporal Integration and Fluctuation Detection.

Despite the highly irregular firing that can be observed in the LIF model through the inclusion of partial reset, it is not clear which are the biological correlates that support his inclusion. In our model, we will include and reinterpret the partial reset mechanism from a metabolic perspective. Specifically, we include the relationship formalized in Eq 5 between ATP level, sodium-potassium pump, and repolarization level through making partial reset mechanism energy-dependent. Our model include ATP dependencies explicitly in *β* := *β*(*A*(*t*)) and follow the rationale explain above through the formalization exhibited on Eq 6:

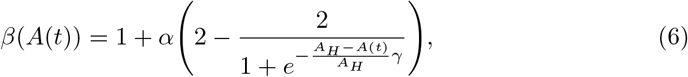

where 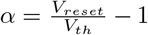. As it can be seen from Eq 6, we are assuming that *β* dependencies on ATP follow a sigmoidal relationship. Using this formalism enables us to represent physiologically plausible repolarization membrane voltage values after an action potential occurs as a function of available ATP in the neuron. Given that the precise curve representing the repolarization membrane voltage dependence on ATP is unknown, we introduce a sensibility parameter *γ*, accounting for this uncertainty. This parameter gives the flexibility to adjust the intensity on which the repolarization membrane voltage value is affected by ATP level changes. To illustrate this relation Fig 1 shows how higher values of the sensibility parameter *γ* account for relationships where repolarization membrane voltage is extremely sensible to ATP displacement from homeostatic ATP (*ATP_H_*), whereas lower γ values represent a smoother transition of *V_reset_* as ATP level changes.

**Fig 1.**
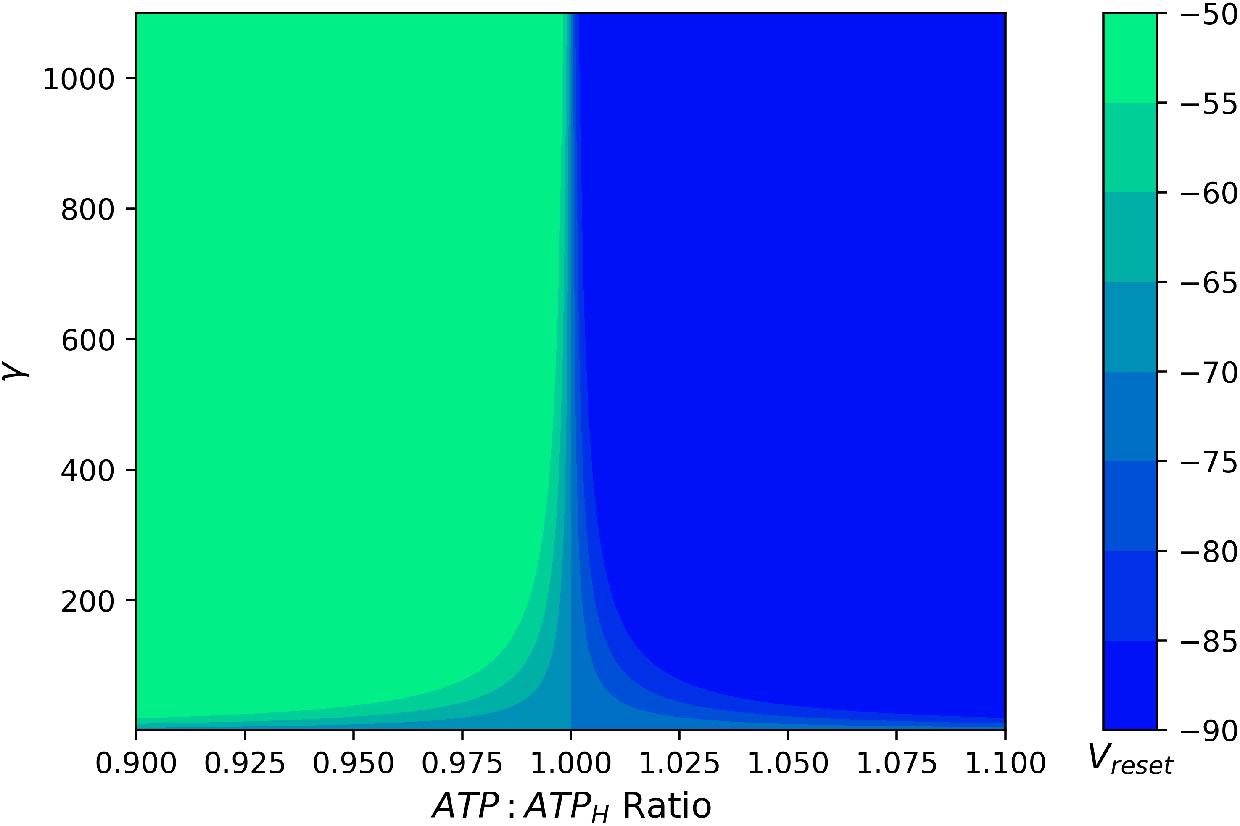
Available ATP and repolarization. Relationship between available ATP ratio (ATP: *ATP_H_*) and reset voltage *V_reset_*, depending on the sensitivity parameter *γ*.

### EDLIF evaluation

To test the EDLIF, we used three strategies:

1. Evaluation of the model’s performance predicting real spike train.
2. Contrasting EDLIF energetic indicated dynamic against actual results.
3. Evaluating if EDLIF was able to reproduce the results of more complex models simulating amyotrophic lateral sclerosis.

For the first strategy, we use publicly available data from *The quantitative single-neuron modeling competition* [3,4] and utilize the *spike-coincidence* (Γ) as a fitness function to measure the performance of each model, allowing to quantify the similarity between real spikes trains and the ones generated by each model. It is also possible to characterize the neuron’s spiking behavior through the *Inter-Spikes-Interval* (ISI) distribution. Fig 2 shows this distribution for the EDLIF and LIF model, and the one generated by a particular trial of neuronal recordings. To quantify the discrepancy between the real ISI distribution and the one generated by the model, the Jensen-Shannon metric [30] is used (Jensen-Shannon metric is bounded between 0 and 1, and is equal to 0 if only the two distributions under comparison are equal).

**Fig 2.**
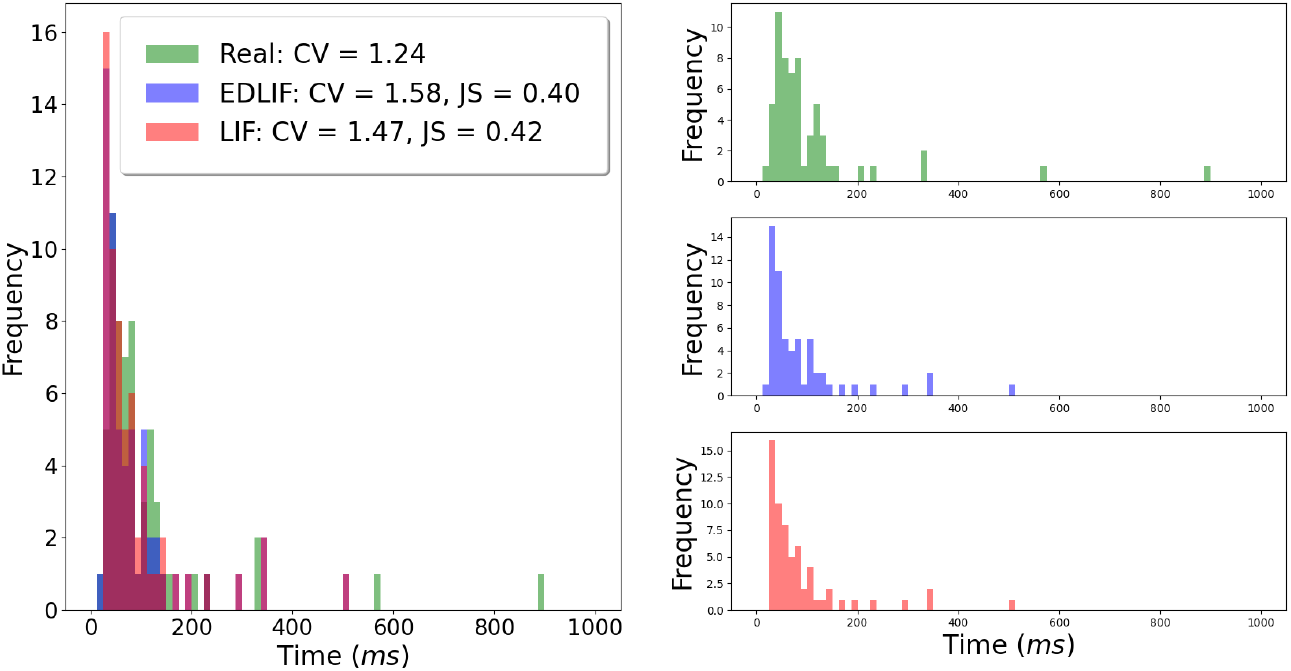
Inter-spike-Interval distributions. Inter-spike-Interval distribution of LIF, EDLIF and real neuron recording. *CV* corresponds to the coefficient of variation, whereas *JS* is the Jensen-Shannon metric, which enables to contrast the discrepancy between the real ISI and the one given by each model. The distribution corresponds to an histogram with 80 bins.

For the second strategy, we evaluated how EDLIF behaves replicating two scenarios, one repeating ATP on-demand production [17,36], and another contrasting EDLIF association of average firing rate and ATP consumption against a thalamocortical biophysically-realistic model [34].

For the third strategy, we explored the predicted effect of EDLIF under a mitochondrial dysfunction scenario. To mimic a mitochondrial dysfunction, we simulated our model considering different levels of reduced homeostatic ATP concentration (*A_H_*) similar to the work of Le Masson and colleagues [7]. Given this scenario, we simulated neuronal behavior with a constant current (4 *seconds*) tuned so that neuron fires at ~ 15 *Hz.*

## Results

We extended the LIF model to include ATP consumption and production in association to neuron physiology. As mentioned above, we will evaluate the extended LIF model (EDLIF) by means of its ability to predict real data, and also to predict a neurodegeneration pattern previously reported.

### Model’s performance predicting real spike trains

In Table 2, we can observe that the EDLIF model has a smaller Jensen-Shannon metric (Column JS metric in Table 2) than the LIF model, implying that the EDLIF model is more akin to the real neuron ISI distribution than the one generated by the LIF model. Surprisingly, although LIF model have smaller refractory time than EDLIF (*τ_r_* in Table 2), in Fig 2 it can be seen that for the EDLIF model there exist ISI’s values in the range (12.5, 25) ms, whereas inter-spike-intervals for LIF model are always above 25 ms. This result can be explained through the energetic dependence introduced to the model through the incomplete repolarization mechanism.

**Table 2.** Model Parameters and Performance. Comparison of LIF and EDLIF model performance in test set, in terms of *spike-coincidence* measure (Γ). JS metric is the Jensen-Shannon metric calculate between experimental ISI distribution and the one generate by the model. *Spike-coincidence* and Jensen-Shannon metric result are average across all trial of neuronal recording. Parameters are obtained by maximizing Γ in train set through PSO optimization algorithm.

| | $C_m$ $nF$ | $V_{th}$ $mV$ | $\tau_r$ $ms$ | $\gamma$ | Performance ($\Gamma$) | JS metric |
| --- | --- | --- | --- | --- | --- | --- |
| LIF | 279.94 | -52.49 | 14.1 |  | 0.55 | 0.41 |
| EDLIF | 202.43 | -51.6 | 16.33 | 178.84 | 0.61 | 0.39 |

Table 2 also presents the parameters (*C_m_, V_th_, τ_r_* and *γ*) when executing the optimization algorithm in each model and the performance (Γ) that each one obtains (reefer to Supporting information for a detailed description of parameter selection through optimization algorithm, dataset used and *spike-coincidence* measure).

These results show that EDLIF have better performance than LIF model, in the sense that it gives a better characterization of real neuron spikes trains under intracellular current stimulation, either considering the *spike-coincidence* measure (Γ), or by contrasting the similarity between model and real ISI distribution by means of Jenssen-Shannon metric.

### Neurons energetics and behavior

To explore the relationship between energetics and neuronal behavior, we submitted the EDLIF model to different experiments. Fig 3 shows how stimulating the neuron affects ATP dynamics. When pulses of 600 *pA* amplitude and 10 seconds width were applied, the available ATP in the neuron starts to drop. These stimuli generate a firing rate of approximately 36 *Hz* and cause a maximum of ~ 0.4% decrease in available ATP. Those slightly ATP decreases mentioned earlier may be explained by the tight coupling between ATP consumption and production. Thus, showing that ATP consumption is rapidly supplied by ATP production, allowing the neuron to maintain and stable ATP concentration. These results are in line with previous studies that show the neuronal capacity to increase ATP production ‘on-demand’ while consumption is increased [17,36].

**Fig 3.**
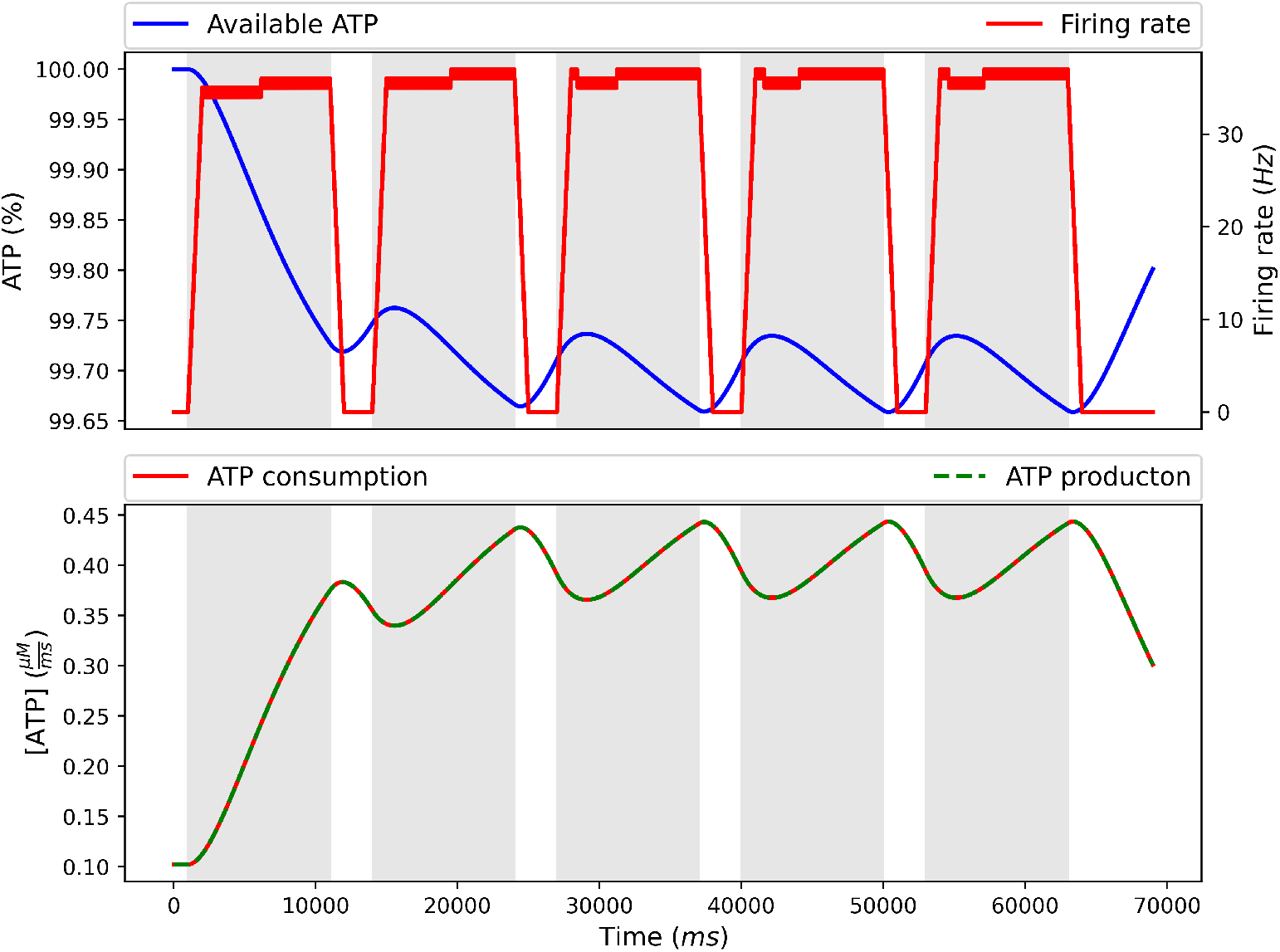
Neuronal behavior and ATP dynamics. Neuronal behavior and ATP dynamics under 60 second stimuli: each pulse have an amplitude of 600 *pA* and 10 seconds width with a 3 *sec* 0 *pA* amplitud between each. Available ATP is measured in percentage with respect to homeostatic ATP level *A_H_* (100% level means that available ATP is equal to *A_H_*).

Regarding the relationship between average firing rate and ATP consumption, in [34] a thalamocortical biophysically-realistic model is used to show that average firing rate, rather than temporal pattern, determined the ATP cost across firing patterns. To study the relation between the average firing rate and ATP consumption, the EDLIF model has been tested under different firing rates. Fig 4 presents that ATP demand increased linearly as a function of firing rate in concordance with [34]. The increase in ATP consumption following raise in firing rate, could be explained by the depolarization phase of action potential generating abundant *N a*^+^ entry, which dominated the metabolic cost of neuronal activity [34]. In summary, our model shows that there is a straightforward relation between ATP consumption and neuronal firing rate intensity, but because of the tight coupling between ATP consumption and production, the available ATP changes slightly.

**Fig 4.**
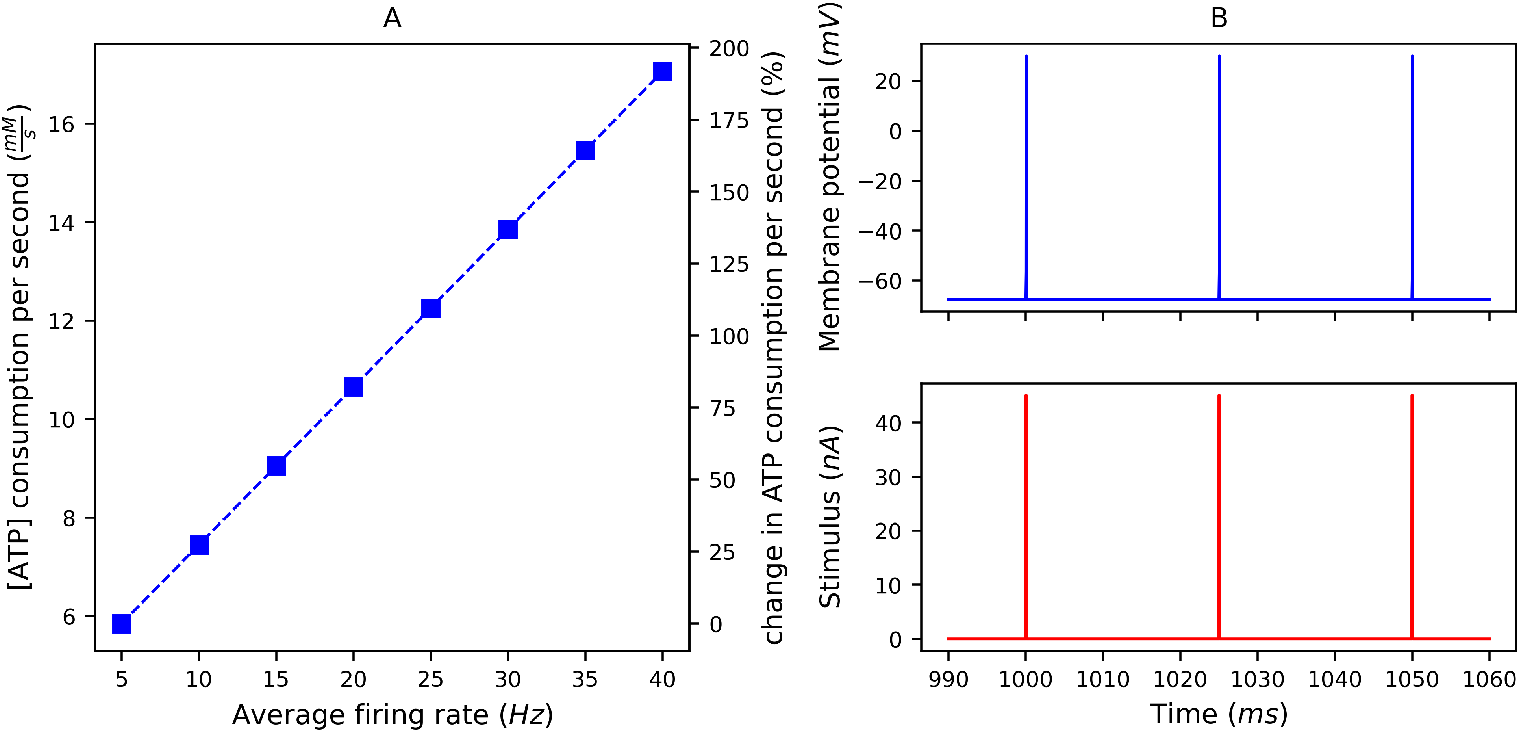
Effect of average firing rate on estimated metabolic cost. (A) Total and relative (%) change in [ATP]/second consumption for different firing rates. (B) neuron’s stimuli and corresponding membrane potential responds (*mV*). The stimuli consisted of 30 seconds of pulses of 0.1 *ms* width and 45 *nA* amplitude.

### Neurodegeneration an energetics: amyotrophic lateral sclerosis

Because our model accounts for energy-dependencies affecting neuronal activity, it should be useful for studying neuronal behavior under a metabolic disorder. Specifically, to explore the link between bioenergetics and neuron degeneration, we use the EDLIF model and simulate neuronal behavior by stimulating the neuron with constant current (4 *seconds*) under mitochondrial dysfunction. As mentioned above, mitochondrial dysfunction was achieved by reduced homeostatic ATP concentration (*A_H_*), following Le Masson and colleagues [7]. As shown in Fig 5, a more substantial mitochondrial dysfunction (lower *A_H_* concentration) implies a higher firing rate response under the same stimuli. This shows how an energy depletion in the neuron affects sodium-pump activity and, consequently, the repolarization process. This entails a depolarization state, which increases the likelihood of an action potential, a hyperexcitable state. The inclusion of the sensibility term (γ in Eq 6) in the EDLIF model accounts for the different affection of energy imbalance in a particular neuron type; therefore, the model is suitable to describe energy-dependent sensitivity dynamics.

**Fig 5.**
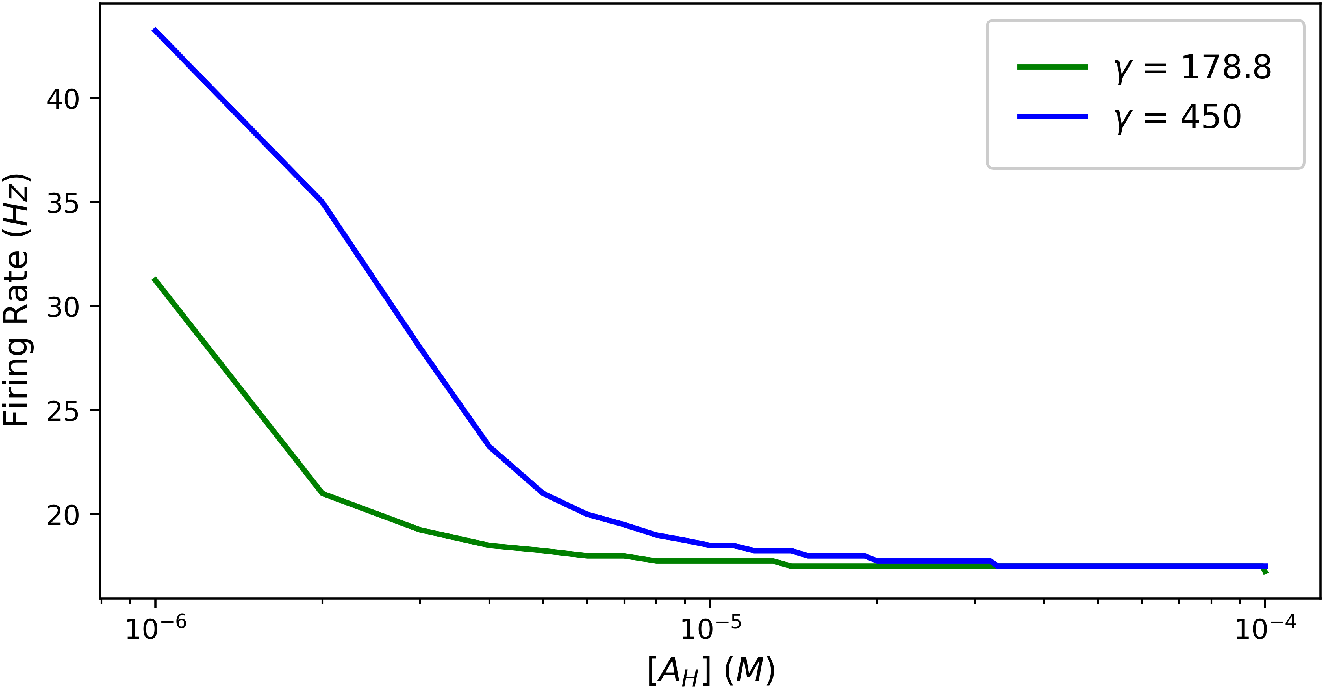
Neuron behavior under ALS. Simulation of ALS disease by reduction of homeostatic ATP level (*A_H_*). As *ATP_H_* is reduced, neurons depolarize and their firing rate increases. When *ATP_H_* is reduced, higher sensitivity values (γ) imply greater firing rate. These results are coherents with the effects of ALS in motor neurons described in [7].

## Discussion

This article has introduced the EDLIF model, an energy-dependent extension of the classical LIF model. Our model aims to integrate the effects of ATP imbalance levels in single-neuron dynamics through partial repolarization mechanisms. The biologically inspired inclusion of ATP-dependence trough partial reset mechanism allows the model to keep computational simplicity and improve predicting capabilities when representing real spike trains considering both *spike coincidence* measure and the dissimilarity (*Jensen-Shannon metric*) between the real ISI distribution and the one generated by the models. Also, the EDLIF model can replicate specific ALS neuronal behavior similar to the model introduced in [7], but using a significantly more straightforward computation. In this sense, the EDLIF model seems suitable to study ALS’s effect *in silico* spiking neural networks with thousands of neurons. It has also been quantified how the average firing rate affects ATP consumption, indicating that the action potential’s firing rate makes a significant contribution to overall energy consumption. These results are essential for interpreting the signals from metabolism-dependent modalities of functional brain imaging [34].

Despite the advantages that the model presents concerning LIF and other more complex models, it is worth to mention some limitations that it offers. Firstly, given that our goal is to develop a simple model that can simulate networks of thousands of neurons, the EDLIF model neglects calcium kinetics and morphological or spatial description, *i.e.* EDLIF is a single-compartment model. The EDLIF model presented here has neglected adaptive functions like the ones shown in the adaptive LIF model or Izhikevich model. Adaptive functions as the one presented in adaptive LIF could be included easily, but it is out of the scope of this work. Evidence suggests that the sodium-potassium pump plays a significant role in neuronal activity regulating membrane potentials and neuronal firing. Experiments show that pharmacological blocking of sodium-potassium pumps increases the spontaneous firing rate and generates membrane depolarization [32]. It has also been shown that there is a membrane voltage depolarization of the neurons during hypoxia and ischemia. This increase is due in part to the inhibition of the Na-K pump due to lowered ATP levels [33]. These experiments emphasize the relevance of Na-K pumps in neuronal activity and their relationship with ATP levels. In this regard, the EDLIF model includes an ATP imbalance in the Na-K pump through incomplete repolarization mechanisms. Thus, making the model a suitable tool for studying the effect of ATP imbalance in neuronal behavior through the accounting of the sodium-potassium pump inhibition when ATP level is low -as in hypoxia- and its consequences in neural and network activity.

Also, ALS results are coherent with other neurodegenerative conditions as the abnormal high-frequency burst firing in rapid-onset dystonia-parkinsonism, which suggest that partially blocking sodium-potassium pumps in the cerebellar cortex is sufficient to cause dystonia, increasing the firing rate of cells and ultimately caused a conversion from tonic firing to high-frequency bursting activity [31]. Fremont and colleague’s work also shows the relationship between the sodium channel density and sensitivity to a partial block of the sodium-potassium pump. A lower sodium channel density implies less sensitivity to the partial blockade of sodium pumps. Thus, γ can be explained in biological terms associating it to the density of sodium channels and disclosing why different *γ* values better describe different neuron type’s behavior in the EDLIF model. Finally, considering the vast literature linking metabolic dysfunction and neurodegenerative disease [12–17,35], simple computational models as the one introduced in this article become relevant as an alternative to classical LIF because of their capability to include energetic dependencies and so making it suitable to study the effect of metabolic disorder from a -*in silico*- networks perspective.

## Acknowledgments

This work has been supported by the supercomputing infrastructure of the NLHPC (ECM-02) a Grant ICM ICN09_015 tp P.E.M The authors also want to thank ANID-PFCHA/Doctorado Nacional/2019-21190330 for supporting Ismael Jaras doctoral studies.

## Author Contributions

IJ and TH conceived and designed the experiments, performed the experiments and analyzed the data. IJ, TH, MEO, PEM, RCV wrote the paper.

## Supporting information

### Determining *τ_AP_*

To determine parameter *τ_AP_*, a detailed biophysical model of the brain’s metabolic interactions is used. The model integrates three modeling approaches; the Buxton-Wang model of vascular dynamics, the Hodgkin-Huxley formulation of neuronal membrane excitability, and a biophysical model of metabolic pathways [26]. To reach equilibrium, the model is left without stimulation for more than 300 seconds. Then, the neuron is stimulated to only produce one spike (Fig 6) and study the ATP dynamic imbalance that it produces (Fig 7).

**Fig 6.**
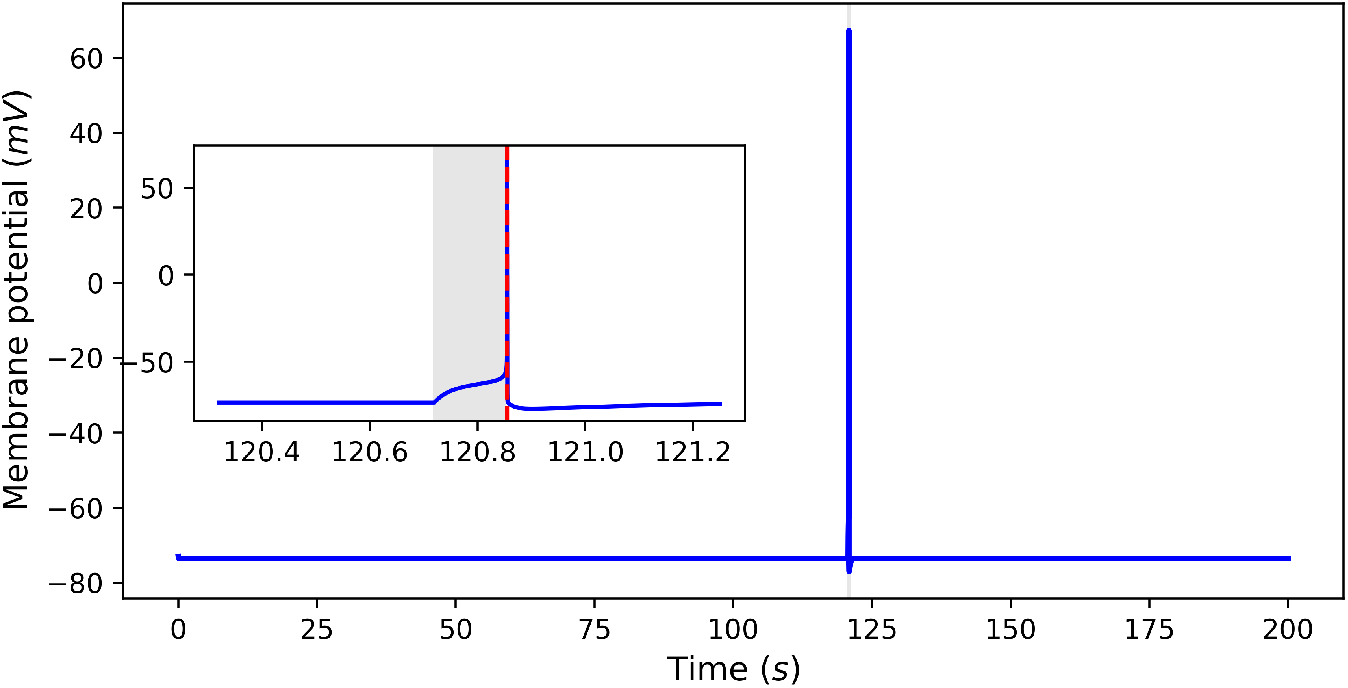
Metabolic-dependent Hodkin-Huxley membrane potential. Membrane potential generated by a detailed Hodkin-Huxley model accounting for biophysical description of metabolic pathways [26].

**Fig 7.**
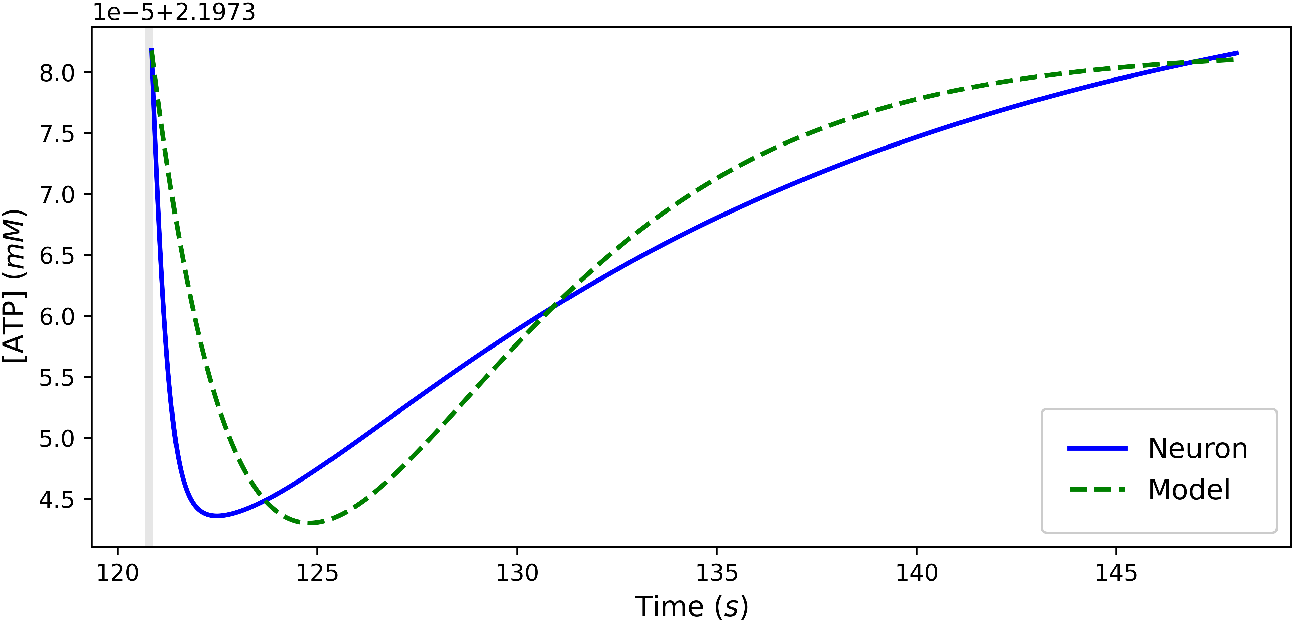
Available ATP dynamic and spike occurrence. The occurrence of one spike produces the ATP dynamic shown in the blue line, while the green line shows the model’s curve obtained by fitting *τ_AP_* in Eq 4 through least square minimization.

### Parameter Fitting

To fit our model to real spike trains and compare performances between EDLIF and LIF, we optimized both models using the dataset supplied by QSNMC (‘‘Quantitative Single Neuron Modeling: Competition 2009” [3,4]). QSNMC data consists of 13 voltage recordings of a layer five pyramidal neuron with the same stimulation for 39 sec. Data were divided into training and test sets. Firstly, we fitted membrane conductance (*g_L_* = 32.9 μΩ^-1^), resting membrane potential (*V_reset_* = –67.54 *mV*), and equilibrium potential of leak channel (*E_L_* = —67.54 *mV*) from the resting state. Specifically, we obtained them from the first five *sec* of voltage recordings of the QSNMC dataset. We used PSO with a split train dataset for the rest of LIF and EDLIF models’ parameters. We used first 80 % recordings from 5 *sec* for training, and the rest 20 % for the test.

We used the PSO algorithm for optimization, which was inspired by birds’ flocking behavior where individuals are sharing the best positions for getting food and finally converge to that point [2]. This algorithm seeks the best position of particles where the cost is minimum or benefit is maximum. For the benefit function, we measured spike coincidence Γ (Eq 7) between two spike trains based on the competition method [3,4]:

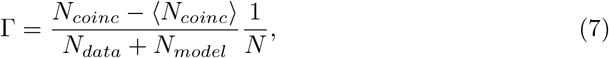

where *N_data_* is the number of spikes on real neuron recording whereas *N_model_* the number of spikes given by the model, one spike coincidence is counted if the spike of the model exists within four *ms* (precision Δ) from that of reference. *N_coinc_* is the total number of spike coincidences and (*N_coinc_*) = 2*v*Δ*N_data_* is the expected number of spike coincidences generated by an homogeneous Poisson process with the same frequency (*v*) as the spike trains associated to *N_model_*. Finally *N* = 1 – 2*v*Δ is a normalization term. To consider and measure spike trains’ reliability, we calculated the intrinsic reliability of reference spike trains. For a detailed procedure to calculate intrinsic reliability and more information about spike coincidence, the reader is referred to QSNMC articles [3,4].

Finally, PSO seeks the best position around a bounded space, so we set upper and lower bounds of each parameter. The bounds are presented in Table 3.

**Table 3.**
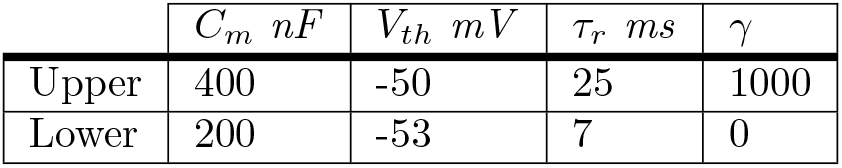
Upper and lower bounds of parameters. Upper and lower bounds of parameters for optimization procedure. PSO algorithm fitted *C_m_*, *V_th_* and *τ_r_* for LIF, and *C_m_*, *V_th_*, *τ_r_* and γ for EDLIF.

## Code availability

Code available on request from the authors.

## Data availability

Data available at: https://github.com/INCF/QSNMC.

